# Extraction of biological terms using large language models enhances the usability of metadata in the BioSample database

**DOI:** 10.1101/2025.02.17.638570

**Authors:** Shuya Ikeda, Zhaonan Zou, Hidemasa Bono, Yuki Moriya, Shuichi Kawashima, Toshiaki Katayama, Shinya Oki, Tazro Ohta

**Affiliations:** Database Center for Life Science, Joint Support-Center for Data Science Research, Research Organization of Information and Systems; Graduate School of Integrated Sciences for Life, Hiroshima University; Institute of Resource Development and Analysis, Kumamoto University; Genome Editing Innovation Center, Hiroshima University; Department of Artificial Intelligence Medicine, Graduate School of Medicine, Chiba University; Institute for Advanced Academic Research, Chiba University

## Abstract

BioSample is a comprehensive repository of experimental sample metadata, playing a crucial role in providing a comprehensive archive and enabling experiment searches regardless of type. However, the difficulty in comprehensively defining the rules for describing metadata and limited user awareness of best practices for metadata have resulted in substantial variability depending on the submitter. This inconsistency poses significant challenges to the findability and reusability of the data. Given the vast scale of BioSample, which hosts over 40 million records, manual curation is impractical. Rule-based automatic ontology mapping methods have been proposed to address this issue, but their effectiveness is limited by the heterogeneity of BioSample metadata. Recently, large language models (LLMs) have gained attention in natural language processing and have been expected as promising tools for automating metadata curation. In this study, we evaluated the performance of LLMs in extracting cell line names from BioSample descriptions using a gold-standard dataset derived from ChIP-Atlas, a secondary database of epigenomics experiment data, which manually curates samples. Our results demonstrated that LLM-assisted methods outperformed traditional approaches, achieving higher accuracy and coverage. We further extended this approach to extraction of information about experimentally manipulated genes from metadata where manual curation had not yet been applied in ChIP-Atlas. This also yielded successful results for the usage of the database, which facilitates more precise filtering of data and prevents misinterpretation caused by inclusion of unintended data. These findings underscore the potential of LLMs to improve the findability and reusability of experimental data in general, significantly reducing user workload and enabling more effective scientific data management.

## Introduction

In recent years, advances in technologies such as high-throughput sequencing for analyzing nucleotide sequences have generated vast amounts of experimental data in the life sciences. To share and publish such experimental data, various public data repositories have been developed, such as the Sequence Read Archive (SRA) [1] for nucleotide sequence data and the Gene Expression Omnibus (GEO) [2] for gene expression analysis data. The number of experiments submitted to these repositories has been increasing, and as of November 2024, there are over 620,000 projects in SRA and 240,000 projects in GEO. Secondary analysis of accumulated public data enhances the reliability of experimental results in subsequent studies and provides additional biological insights beyond those obtained by the original submitters. Moreover, services called secondary databases have been developed to collect public data on specific experiment types and provide interfaces for browsing and analyzing such data. Examples include DEE2 [3] and GREIN [4] for gene expression analysis, and ChIP-Atlas [5] for epigenomics analysis such as Chromatin Immunoprecipitation followed by sequencing (ChIP-seq).

Users seeking data of interest from these repositories with massive amounts of data typically search based on experimental conditions and sample information. The information describing “data about experimental data”, is referred to as metadata. Initially, sample metadata was recorded in sample-specific records for each data repository. However, as more analyses are conducted on the same samples, managing and searching for metadata within individual repositories has become increasingly cumbersome. To address this, the BioSample database was developed to centrally store sample information independent of experimental type [6]. As of November 2024, BioSample hosts over 40 million records.

In BioSample, metadata such as organism, tissue or cell type, disease, and treatment conditions is described as key-value pairs (Fig. 1). The experimental conditions that should be described as metadata vary widely, making it difficult for the database to define standardized rules to describe them. While packages specifying the required metadata for certain experiment types have been introduced, a Generic package also exists with no predefined requirements. According to Gonçalves and Musen [7], 85% of BioSample records use the Generic package. Consequently, much of the metadata description is left to the discretion of submitters, resulting in inconsistencies even for identical experimental conditions. This situation undermines the purpose of public data repositories, which is to enable data reuse by other researchers.

**Fig. 1.**
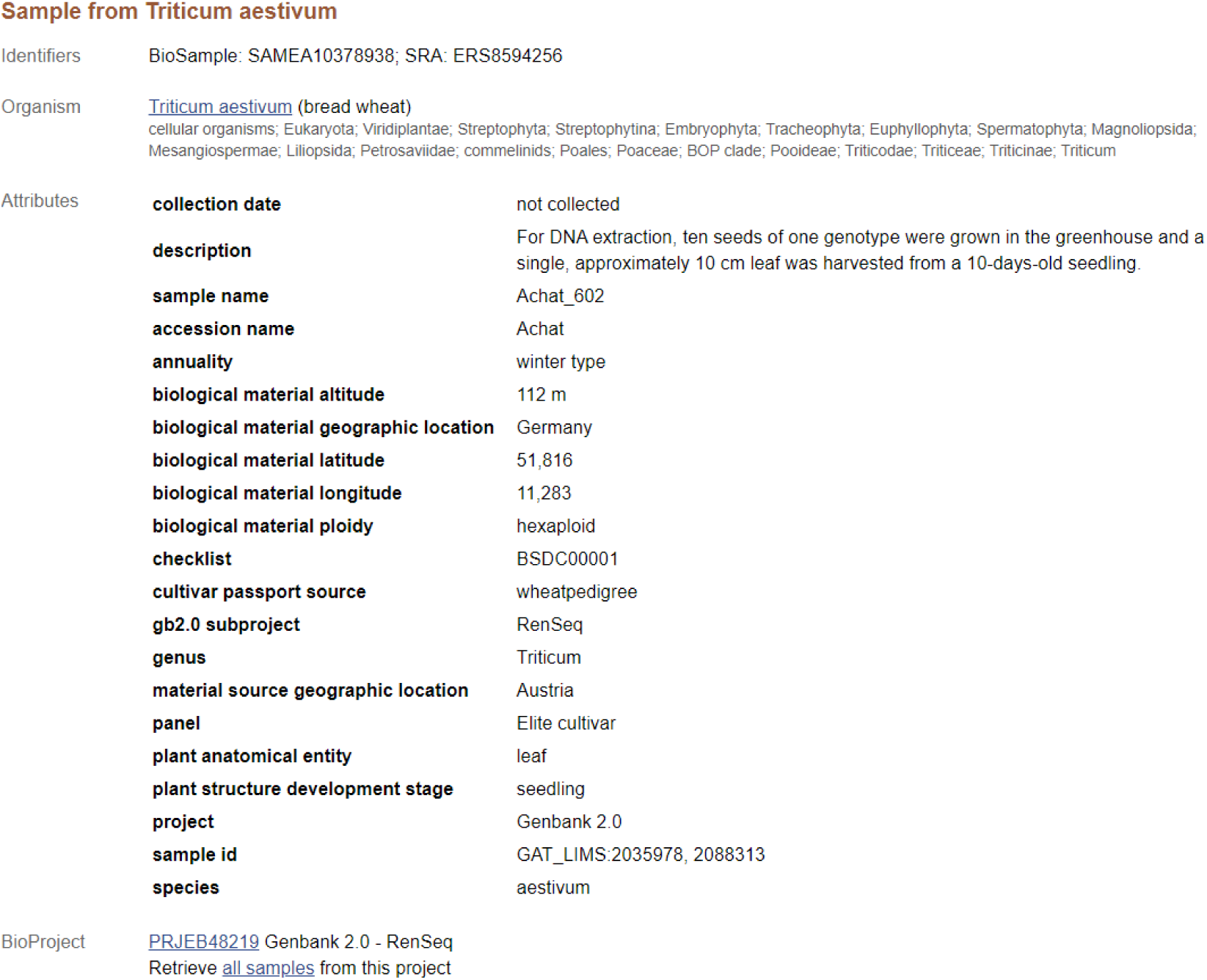
An example of BioSample records (https://www.ncbi.nlm.nih.gov/biosample/SAMEA10378938). The attributes of the sample are described as a set of pairs of attribute keys like “plant anatomical entity” and their corresponding values like “leaf”.

One issue is the use of multiple representations for the same concept. For instance, synonyms (e.g., neuron vs. nerve cell), abbreviations and full names (e.g., hESC vs. human embryonic stem cell), variations in capitalization, and typographical errors caused by human error are common. Users of BioSample face difficulties retrieving all samples of their interest, because there is no unified terminology for describing concepts. To resolve this, using ontologies to describe metadata in BioSample submissions can be helpful. Ontologies structure domain-specific concepts semantically by defining hierarchies and synonyms for terms. Each component of ontologies, called as ontology terms, standardizes the description of concepts. Examples of ontologies in the life sciences include UBERON [8] for anatomical concepts, Cell Ontology [9] for cell types, and Disease Ontology [10] for human diseases. If BioSample metadata were described using ontology terms, it would alleviate the difficulty of retrieving samples with identical conditions. However, ontology usage in BioSample remains limited. For example, while BioSample packages define that the “disease” attribute should use Disease Ontology terms for human samples (https://www.ncbi.nlm.nih.gov/biosample/docs/attributes/), an investigation in November 2024 revealed that only 148,876 out of 595,177 human samples with a “disease” attribute used strings matching Disease Ontology labels.

Several strategies can be considered to map BioSample metadata to ontologies. Manual curation by experts is the most primitive approach and can achieve high accuracy, but this suffers from low scalability. In ChIP-Atlas, for example, metadata from epigenomics experiments is manually annotated by experts using a controlled vocabulary. However, this manual curation is limited to specific attributes, such as antigens and cell types, and expanding its scope would require additional effort. Therefore, automated curation systems are needed if scalability is prioritized over accuracy.

An example of efforts to automatically map BioSample metadata to ontologies is MetaSRA [11]. MetaSRA maps key-value pairs in BioSample records to concepts such as tissue, cell type, cell line, disease, and developmental stage, using ontologies like UBERON, Cell Ontology, Cellosaurus [12], Disease Ontology, and Experimental Factor Ontology [13]. MetaSRA employs fuzzy string matching to query ontologies for terms and maps identified terms to metadata. However, this strategy struggles with homonyms and fails to distinguish terms used in a negative context. While MetaSRA applies rules to reduce misannotations—such as permitting mapping to cell line terms only for attributes named “cell line” or “cell type”—information may still be missed since cell line data is not always described under these specific attribute names.

Moreover, inappropriate mappings may persist despite these rules. For instance, when long text descriptions are provided in attribute values, some strings may not represent the sample itself. Fig. 1 illustrates a wheat sample with an attribute named “plant anatomical entity” to indicate this sample derives from a leaf. This record also has the “description” attribute to describe the sampling protocol in natural language. However, the presence of the words “seeds” and “leaf” within this description poses a challenge for rule-based ontology mapping. While humans can easily determine that this sample is collected from a leaf, it is difficult to algorithmically determine that the word “seeds” appears in a procedural explanation and does not represent that this sample is not a seed.

To address the challenges inherent in rule-based approaches, machine learning-based methods have been considered. However, conventional machine learning techniques have struggled to address the vast variety of description patterns in BioSample due to the difficulty of preparing a sufficiently comprehensive training dataset. For instance, Klie et al. aimed to enhance metadata attributes in BioSample using a deep learning-based approach [14]. They tackled a named entity recognition (NER) task, extracting word sequences from longer text, such as sample titles, that likely represented values for specified attributes. They used key-value pairs of BioSample as training data to develop the model to learn strings deemed plausible as values for given attributes. While the model achieved high accuracy in extracting strings, maintaining this level of accuracy required excluding monograms from the training set, which posed a limitation for extracting concepts represented by single-word terms. Furthermore, the approach relied on straightforward extraction and lacked the ability to differentiate between strings used in negative contexts or other nuanced scenarios.

Recent advances in large language models (LLMs) trained on diverse, extensive datasets have outperformed traditional methods in natural language processing tasks. While BioSample lacks consistent rules to describe metadata, it is typically understandable to humans. LLMs could potentially interpret such metadata and reorganize it appropriately for describing samples. The application of LLMs to NER and organization of academic terminology is actively researched. For instance, Dagdelen et al. [15] demonstrated the use of LLMs to extract specific information from materials science papers and structure it as JSON objects. Similarly, Sundaram et al. [16] reorganized BioSample metadata according to attributes defined in existing metadata support tools. These studies highlight the potential of LLMs to improve metadata organization and searchability.

As discussed, LLMs are expected to be effective to address challenges in BioSample metadata such as inconsistent attribute names and values, as well as the semantic interpretation of text strings, enabling high-accuracy concept extraction that was previously difficult to automate. Applying this approach to concepts not covered by manual curation could enhance secondary databases built on BioSample data, improving user experience through increased searchability. In this study, we first examined the previously noted heterogeneity in BioSample descriptions from a different perspective and validated the feasibility of using LLMs for BioSample metadata curation. Then we evaluated the effectiveness of current LLMs in curating BioSample metadata. To quantitatively assess concept extraction, a gold standard dataset was constructed based on ChIP-Atlas’ manually curated results. Evaluation demonstrated that LLM-based methods outperformed traditional approaches. Furthermore, the application of LLMs to extract experimentally manipulated gene names from metadata was conducted and manually evaluated, showing that LLMs achieved sufficient accuracy to aid users in refining their searches, despite some limitations posed by the complexity of BioSample descriptions.

## Methods

### Construction of the Gold-standard Dataset for Cell Line Name Extraction

To quantitatively evaluate the extraction task by LLM, we constructed a gold-standard dataset which defines ontology terms to represent BioSample records. For its creation, the manual curation results from ChIP-Atlas, an integrated epigenomics database, were utilized. ChIP-Atlas comprehensively collects ChIP-seq, Assay for Transposase-Accessible Chromatin with sequencing (ATAC-seq), and Bisulfite sequencing from SRA without any filtering. ChIP-Atlas manually maps information on the tissues and cell types of sample origins to its own controlled vocabulary, with the expertise of developmental biology specialists. While the curated results from ChIP-Atlas are not mapped to any ontology, leveraging this curated dataset was deemed a more efficient and reliable approach for determining ontology terms representing the BioSample records, compared to building one from scratch.

The following considerations were taken into account when selecting samples: 1) Metadata for samples used in ChIP-seq experiments usually include the names of proteins targeted by ChIP. Protein names, which typically consist of alphanumeric combinations, bear similarity to cell line names, and therefore the presence of protein names in metadata could influence the difficulty of the cell line extraction task. Meanwhile, ATAC-seq experiments do not target specific proteins and do not suffer from this issue. Thus, we selected 300 samples each from ChIP-seq and ATAC-seq experiments and enabled evaluation within each experiment type. 2) To avoid skewing the task’s difficulty, the 300 samples selected from each experiment type were ensured to come from distinct projects. Similarly, samples with identical terms mapped by ChIP-Atlas curation results were excluded. 3) We included only human samples, because of the availability of Cellosaurus, an ontology that includes over 110,000 human cell lines and enables precise definition of cell line terms representing BioSample records.

For the selected samples, corresponding terms from the Cellosaurus ontology were identified and defined as the gold standard.

### Automated Annotation Using LLM

#### Setup for LLM Execution

We employed Ollama [17] to run the Llama 3.1 70B instruct q4_0 model [18] on a machine equipped with an NVIDIA RTX 6000 Ada GPU (48 GB memory). To ensure reproducibility of results, the temperature parameter was set to 0. The source code for task-specific prompts and input-output processing has been made publicly available on GitHub (https://github.com/sh-ikeda/bsllmner).

#### Cell Line Name Extraction Task

We designed a pipeline for performing the cell line name extraction task (Fig. 2). In the task, we used the set of the attributes describing each BioSample record as input, prompting the LLM to extract the name of a cell line that is considered to represent the sample. The prompt (Prompt 1) provided a general definition of cell lines, followed by instructions to analyze JSON-formatted data that contain key-value pairs of the sample attributes. The LLM was tasked to determine whether the sample was a cell line and, if so, to extract the cell line name. The extracted cell line names were used as the values for the “cell_line” attribute in JSON files, which were then processed using MetaSRA to map them to ontology terms. We did not use LLMs for the ontology mapping due to the frequently observed hallucination issues where irrelevant ontology term IDs are presented, a problem inherent in LLMs. For samples resulting in multiple ontology terms with the same cell line name, further refinement was performed by the LLM: each ontology term’s description was provided to the LLM, which compared this information with the BioSample metadata to output the most appropriate term (Prompt 2). The Cellosaurus information used in this process included the main label of the cell line (“name”), synonyms (“related_synonyms” and “exact_synonyms”), associated diseases (“diseases”), cell line type such as cancer cell line or embryonic stem cell line (“cell line type”), and the sex of the originating individual (“sex”). These details were appended to the end of Prompt 2 in JSON format, as shown below.

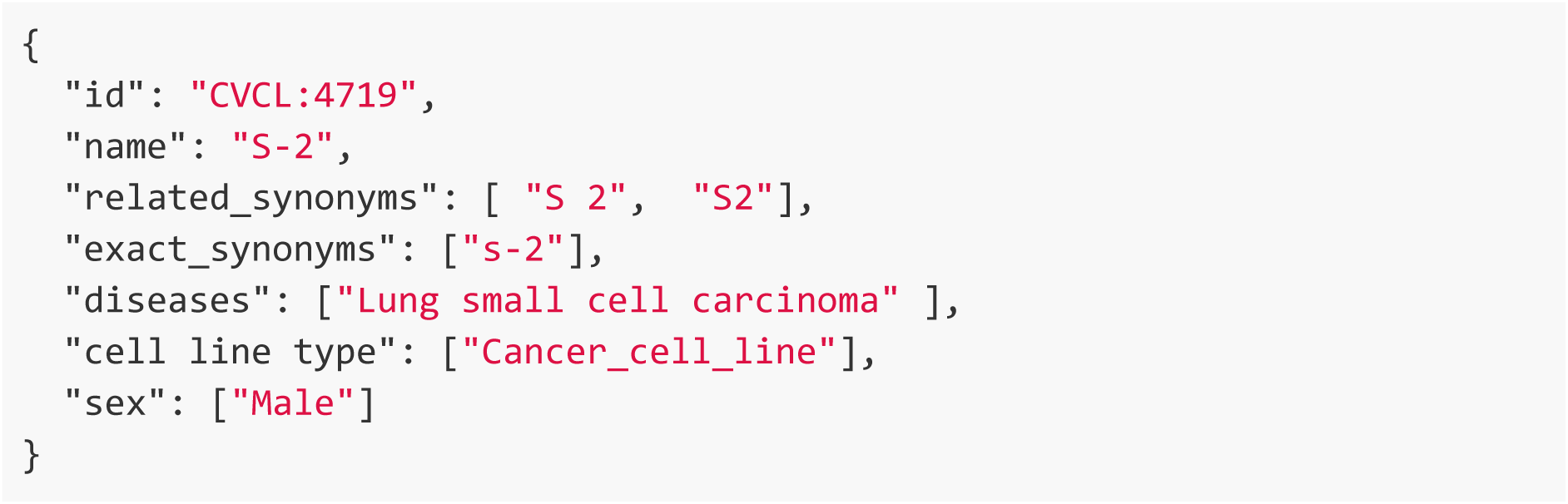

**Fig. 2.**
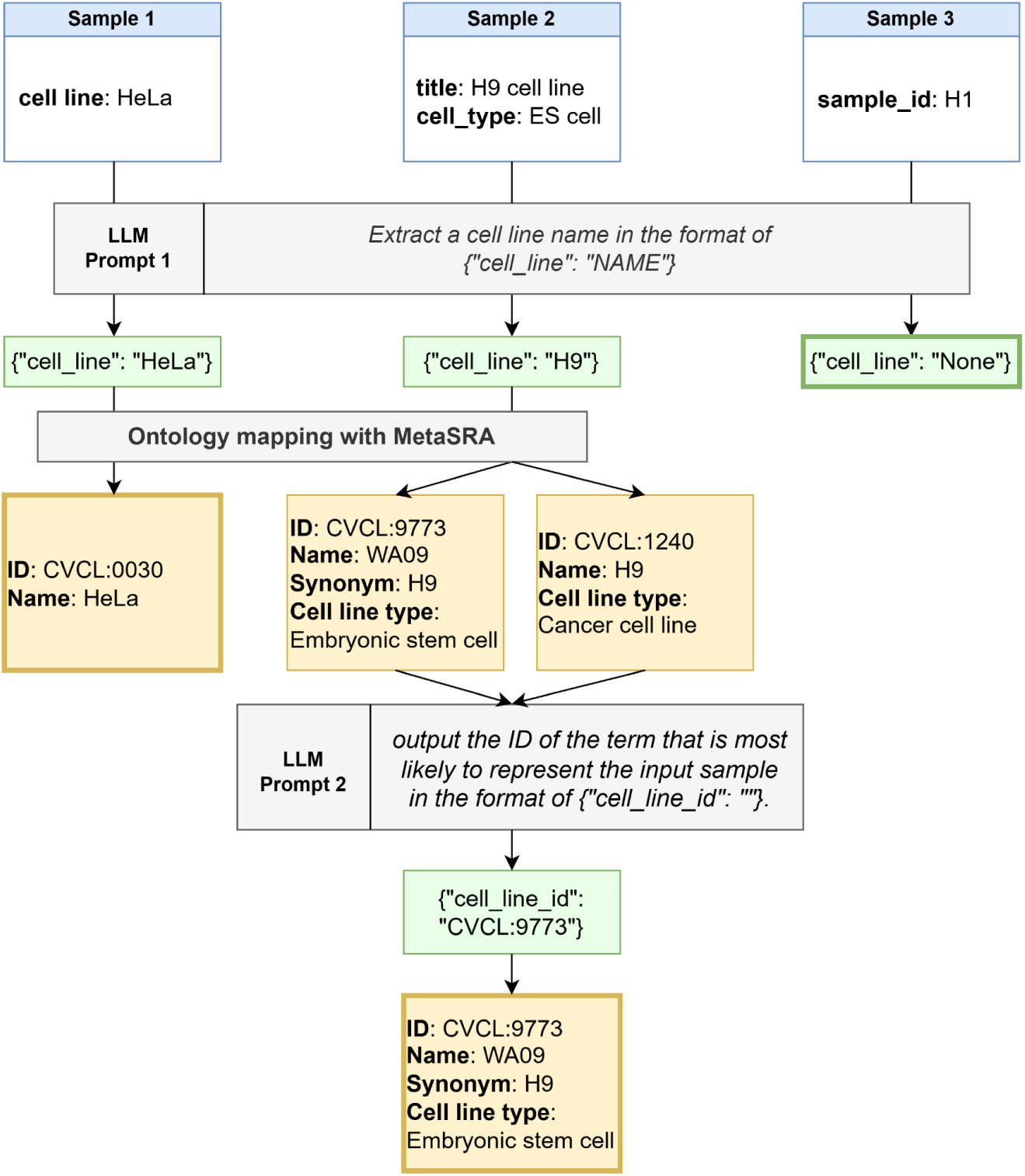
A flowchart describing the LLM-assisted ontology mapping pipeline of BioSample.

To improve performance, the “Think-step-by-step” method was applied in the prompts. This method adds the phrase “Think step by step.” to the end of the prompt, encouraging the LLM to output not only the solution but also the reasoning process, a practice reported to enhance accuracy [19].

#### Ontology Mapping

We used a method based on MetaSRA for ontology mapping from extracted strings to Cellosaurus terms (https://github.com/sh-ikeda/MetaSRA-pipeline). Since the original MetaSRA pipeline (https://github.com/deweylab/MetaSRA-pipeline/) was implemented in Python 2, we ported it to Python 3 for better maintainability. Additionally, we made several refinements:

- Improved handling of cases where ontology term labels or synonyms differed from the query only in capitalization, ensuring complete support for such scenarios.

- Replaced delimiter characters such as “_” and “-” in the input data with spaces and included the resulting strings as part of the query.

In the original implementation, strings with a length of 2 were excluded from the search to prevent mismapping, with a few exceptions. We modified this to allow all such strings without restriction because wrongly mapped terms were expected to be filtered in the selection phase by LLMs (Prompt 2).

#### Evaluation Score Calculation

For the comparative evaluation of the existing method and our proposed LLM-assisted methods, we calculated the following metrics (Fig. 3):

**Fig. 3.**
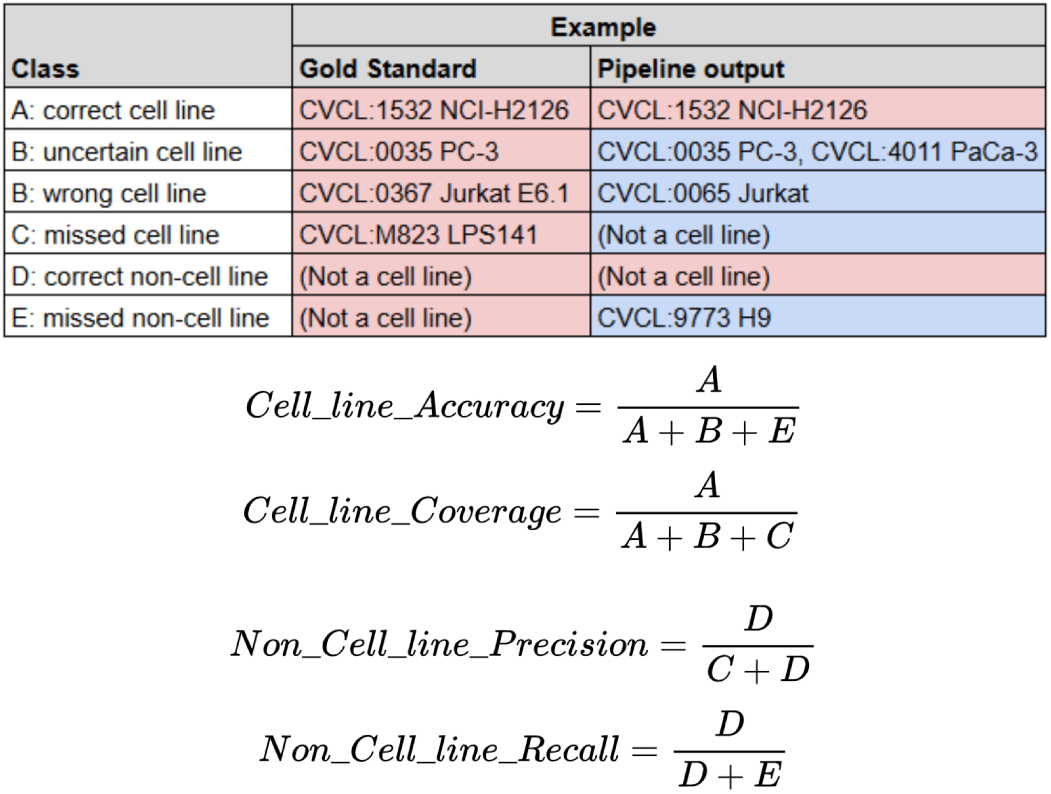
Definition of the metrics used for the ontology mapping pipelines.

- Accuracy and coverage for the task of mapping the correct cell line name to cell line samples.

- **Cell line accuracy** = (Number of outputs mapping to a cell line that are correct) / (Total number of outputs mapping to a cell line)

- **Cell line coverage** = (Number of correct cell line mappings included in the outputs) / (Total number of gold standard entries for cell lines)

- Precision and recall for the task of identifying samples that are not cell lines.

- **Non-cell line precision** = (Number of outputs not mapping to a cell line that are correct) / (Total number of outputs not mapping to a cell line)

- **Non-cell line recall** = (Number of correct non-cell line entries included in the outputs) / (Total number of gold standard entries for non-cell lines)

In cases where multiple cell line terms are suggested by the pipelines, we considered the output incorrect as the correct cell line was not uniquely identified, even if the candidates include the correct cell line.

#### Gene Name Extraction Task

For the task of extracting gene names experimentally modulated in expression, we used human samples from ATAC-seq and ChIP-seq projects registered in ChIP-Atlas. For ATAC-seq samples, we randomly selected one sample from each of the 1,794 human projects registered in ChIP-Atlas. For ChIP-seq samples, since the number of registered projects was relatively large at 4,806, we first randomly selected 2,000 projects as a subset and then randomly chose one sample from each. We used the EBI BioSamples API to obtain the sample data, but data could not be obtained for 42 ATAC-seq samples and 29 ChIP-seq samples. As a result, we used the 1,752 ATAC-seq and 1,971 ChIP-seq samples for the task.

The prompt (Prompt 3) instructed the LLM to analyze JSON-formatted BioSample metadata and output a list of modulated genes and their respective modulation methods as a JSON array. Since a single sample might involve the modulation of multiple genes, the output array consisted of JSON objects with “gene” and “method” attributes, such as:

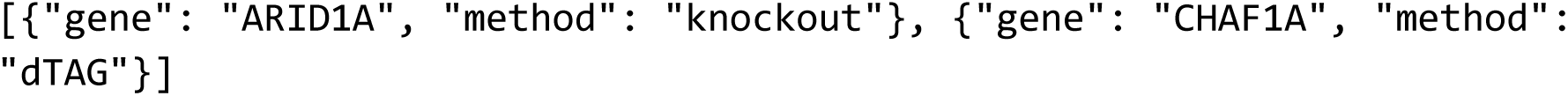

The prompt began by defining the major modulation methods: gene knockout, knockdown, and overexpression. These are prone to significant variability in terminology. For example:

● **Knockout** can appear as “knockout”, “KO”, “-/-”, or “deletion”.
● **Knockdown** may be described using “knockdown”, “KD”, “shRNA”, “siRNA”, “si(Target gene name)”, or “RNAi”.
● **Overexpression** might include terms like “overexpression”, “OE”, “transfection”, or “transduction”.

If a gene was identified in the metadata as modulated by one of these methods, the output’s “method” attribute was aligned to these standard terms. For other methods described in the input, the prompt instructed the LLM to extract and retain the method name as is.

The extracted results were manually evaluated. Samples with descriptions that are difficult to yield a definitive result under the current prompts were excluded from evaluation (examples are given in later sections).

**Prompt 1: Cell line extraction**

> *A cell line is a group of cells that are genetically identical and have been cultured in a laboratory setting. For example, HeLa, Jurkat, HEK293, etc. are names of commonly used cell lines.*

> *I will input json formatted metadata of a sample for a biological experiment. If the sample is considered to be a cell line, extract the cell line name from the input data.*

> *Your output must be JSON format, like {”cell_line”: “NAME”} . “NAME” is just a place holder. Replace this with a string you extract.*

> *When input sample data is not of a cell line, you are not supposed to extract any text from input. If you can not find a cell line name in input, your output is like {”cell_line”: “None”} . Are you ready?*

**Prompt 2: Cell line selection**

> *I searched an ontology for the cell line, “{{cell_line}}”. I have found multiple terms which may represent the sample. Below are the annotations for each term. For each term, compare it with the input JSON of the sample and show your confidence score (a value between 0-1) about to what extent the entry represents the sample. In the comparison, consider the information such as:*

> *Whether the term has a name or a synonym exactly matches the extracted cell line name, “{{cell_line}}”.*

> *Whether the term has disease or cell line type information which matches sample information.*

> *Based on the confidence score, output the ID of the term that is most likely to represent the input sample in the format of {”cell_line_id”: “”}. If it is not clear which one is most likely from the given information, output {”cell_line_id”: “not unique”}.*

**Prompt 3: Gene name and gene modulation method extraction**

> *There are several experimental methods to modulate gene expression.*

> *Gene knockout (KO), also known as gene deletion, involves completely eliminating the expression of a target gene by replacing it with a non-functional version, usually through homologous recombination in cells or animals. This results in a complete loss of the gene’s function.*

> *Meanwhile, gene knockdown (KD), also known as RNA interference (RNAi), involves reducing the expression of a target gene without completely eliminating it. KD is achieved by introducing small RNA molecules, siRNA or shRNA, that specifically bind to and degrade the messenger RNA (mRNA) of the target gene.*

> *Gene overexpression refers to the process of increasing the expression of a specific gene beyond its normal levels in a cell. This is achieved by trasfection of a plasmid carrying the gene of interest, transduction of viruses carrying the gene of interest, etc.*

> *I will input json formatted metadata of a sample for a biological experiment. If the sample is considered to have genes whose expression is experimentally modulated, extract the gene names from the input data and specify the modulation method.*

> *Your output must be in JSON format, like [{”gene”: “GENE_NAME”, “method”: “METHOD_NAME”}]. “GENE_NAME” and “METHOD_NAME” are placeholders. Replace them with the gene name you extract and the modulation method name you specify, respectively. If the modulation method is either gene knockout, gene knockdown, or gene overexpression, the value of the “method” attribute must be “knockout”, “knockdown”, and “overexpression”, respectively. Otherwise, the value of the “method” attribute must be the method name found in the input data.*

> *If the input sample data is not considered to have genes whose expression is modulated, your output JSON must be an empty list (namely,’[]’). Note that multiple genes can be modulated in one sample. In this case, be sure to include all of them in the list of the output JSON. For example, if you find “PRNP” and “MSTN” as knocked out genes, your output must be [{”gene”: “PRNP”, “method”: “knockout”}, {”gene”: “MSTN”, “method”: “knockout”}]. Note also that multiple gene modulation methods can be used for one sample. For example, you may find “ARID1A” as a knocked-out gene and “CHAF1A” as a gene treated with dTAG. In this case, your output must be [{”gene”: “ARID1A”, “method”: “knockout”}, {”gene”: “CHAF1A”, “method”: “dTAG”}].*

> *Are you ready?*

### Mapping gene names to gene IDs

To map gene names extracted using the LLM to gene IDs, we utilized the HGNC multi-symbol checker (https://www.genenames.org/tools/multi-symbol-checker/). This tool allows for searching input strings and retrieving corresponding IDs, including not only Approved official symbols but also Previous symbols, Alias symbols, and Withdrawn symbols. We included all these options for mapping and set the “Search case” parameter to “insensitive”, because gene names in BioSample metadata are not always the current official symbols.

## Results

### Survey of BioSample Metadata

As highlighted by Gonçalves and Musen [7], BioSample attribute names exhibit significant variability, complicating rule-based automated metadata organization. To confirm this, we conducted investigations into several aspects of BioSample metadata variability.

To assess the variability in attribute names used to represent cell lines, we examined the attribute names associated with the value “HEK293T,” a commonly used and distinctive cell line name. It is unlikely for “HEK293T” to refer to anything other than the cell line. Despite this specificity, 27 different attribute names were found to be used for this single cell line name (Table 1), demonstrating significant inconsistency even for identical concepts.

**Table 1.** The variability of the names of attributes whose value is “HEK293T”.

| Attribute name | # of Projects | # of Samples |
| --- | --- | --- |
| cell line | 667 | 11547 |
| cell_line | 435 | 96516 |
| source_name | 338 | 4972 |
| isolate | 88 | 2641 |
| cell type | 33 | 326 |
| strain | 20 | 1104 |
| cell_type | 7 | 507 |
| tissue | 6 | 442 |
| spike-in cell line | 6 | 385 |
| cell line background | 6 | 68 |
| biomaterial_provider | 4 | 102 |
| human cell line | 4 | 23 |
| cell_subtype | 3 | 952 |
| lab_host | 3 | 76 |
| cell line/type | 3 | 14 |
| spike-in cell_line | 2 | 68 |
| cell-type | 2 | 29 |
| host cell line | 2 | 8 |
| cell lines | 1 | 96 |
| mix1 | 1 | 56 |
| cell line/tissue | 1 | 49 |
| cell line/strain | 1 | 26 |
| lab host | 1 | 10 |
| cell line or tissue | 1 | 6 |
| dev_stage | 1 | 4 |
| isolation_source | 1 | 2 |
| cell | 1 | 1 |

We also analyzed the frequency of attribute names associated with values containing the string “H1” (Table 2). “H1” is the name of a commonly used ES cell line, but this simple string can represent a different concept than a cell line. In fact, the attribute names containing “H1” values displayed substantial variability, as shown in Table 2, and included names like “well” and “genotype,” which are unlikely to represent cell lines. This highlights the ambiguity inherent in interpreting such simple strings in metadata.

**Table 2.** The variability of the names of attributes whose value includes “H1”.

| Attribute name | # of Projects | # of Samples |
| --- | --- | --- |
| cell line | 299 | 4734 |
| source_name | 234 | 4360 |
| cell type | 74 | 1043 |
| cell_line | 44 | 635 |
| isolate | 32 | 192 |
| well | 17 | 263 |
| tissue | 13 | 77 |
| treatment | 10 | 54 |
| strain | 9 | 160 |
| genotype | 8 | 59 |
| cell_type | 8 | 99 |
| cell line/strain | 8 | 51 |
| sample type | 5 | 30 |
| sample name | 5 | 25 |
| individual | 5 | 29 |
| Submitter Id | 5 | 5 |
| submitted subject id | 3 | 18 |
| subject | 3 | 128 |
| cellline | 3 | 52 |
| subject id | 2 | 8 |
| sample_patient | 2 | 2 |
| line | 2 | 95 |
| genotype/variation | 2 | 6 |
| es cell line | 2 | 32 |
| donorid | 2 | 187 |
| donor id | 2 | 19 |
| donor cell line | 2 | 116 |
| description | 2 | 14 |
| condition | 2 | 28 |
| cell description | 2 | 9 |
| antibody targetdescription | 2 | 2 |
| LINE | 2 | 95 |
| well position | 1 | 2 |
| well id | 1 | 1 |
| uniqueid | 1 | 1 |
| treatment/time period | 1 | 3 |
| tissue-type | 1 | 1 |
| time | 1 | 12 |
| submitted sample id | 1 | 1 |
| subclone | 1 | 10 |
| stimulus | 1 | 2 |
| state | 1 | 2 |
| source cell line | 1 | 4 |
| source | 1 | 2 |
| short_name | 1 | 1 |
| shRNA | 1 | 2 |
| seq_id | 1 | 2 |
| sample_type | 1 | 3 |
| sample name in supplementary file | 1 | 1 |
| sample description | 1 | 2 |
| replicate | 1 | 8 |
| psc line | 1 | 3 |
| position in smart-seq2 | 1 | 14 |
| position in library | 1 | 11 |
| plate_location | 1 | 5 |
| plate-position | 1 | 4 |
| phenotype | 1 | 1 |
| patient_id | 1 | 1 |
| patient id | 1 | 4 |
| parental cell line | 1 | 8 |
| other sample name | 1 | 1 |
| originating cell line | 1 | 18 |
| original_Sample_ID | 1 | 1 |
| origen of cultured cells | 1 | 4 |
| name in the manuscript | 1 | 24 |
| library | 1 | 3 |
| label | 1 | 2 |
| input | 1 | 44 |
| halplogroup | 1 | 4 |
| growth condition | 1 | 3 |
| grna | 1 | 6 |
| genetic perturbation | 1 | 37 |
| expression | 1 | 3 |
| donor samples | 1 | 1 |
| donor | 1 | 379 |
| differentiated from | 1 | 5 |
| depletion | 1 | 2 |
| crispr library | 1 | 22 |
| column in countmatrix | 1 | 1 |
| clone_id | 1 | 1 |
| cell source | 1 | 8 |
| cell lines | 1 | 576 |
| cell line of origin | 1 | 20 |
| cell line name | 1 | 18 |
| cell line background | 1 | 6 |
| biospecimen repository sample id | 1 | 1 |
| assay name | 1 | 3 |
| antibody | 1 | 9 |
| Sample Name | 1 | 25 |
| Name | 1 | 1 |
| DONOR_ID | 1 | 1 |
| DIFFERENTIATION_METHOD | 1 | 26 |
| ArrayExpress-StrainOrLine | 1 | 9 |

We surveyed the usage frequency of all attribute names. Since samples from the same project often share the same attributes and the number of samples per project is highly skewed [11], we conducted the counting on a project basis. As of June 7, 2024, BioSample contained 27,639,806 records associated with BioProject [6], featuring 76,282 unique attribute names. Of these, 57,750 (75.7%) attribute names were used in only a single project, and 73,207 names (96.0%) were used in 10 or fewer projects.

Examining individual records revealed instances where data submitters seemed unfamiliar with proper metadata annotation practices. Some records contained incomprehensible strings or poorly described attributes, as demonstrated in Table 3 and Supplementary Fig. 1.

**Table 3.**
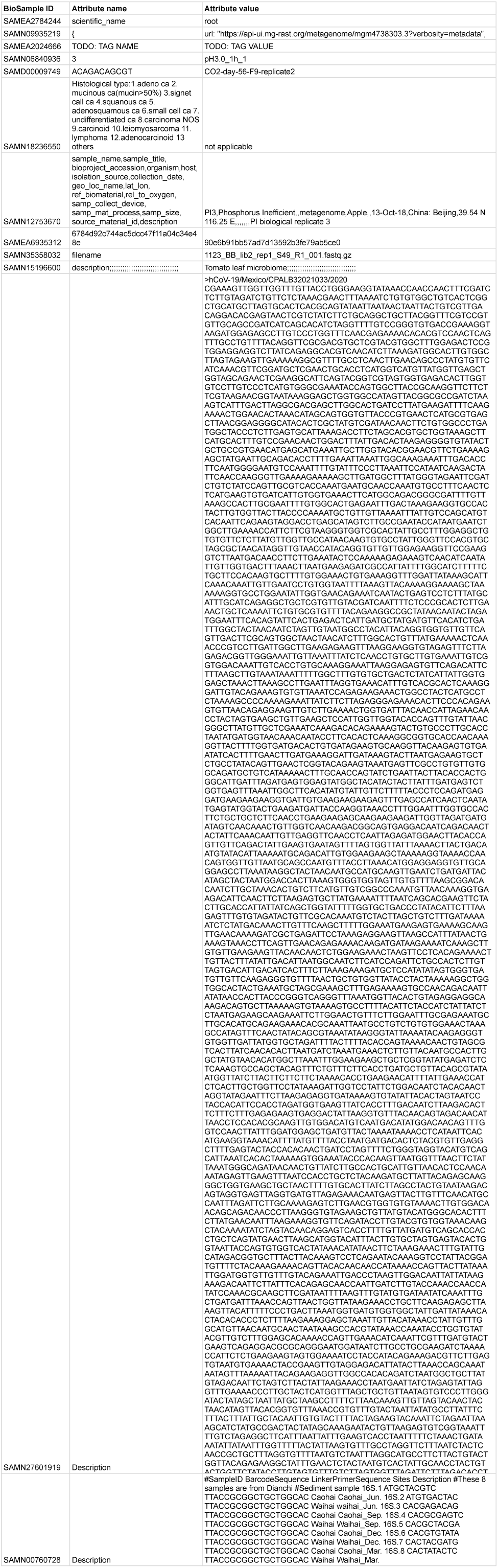
Examples of sample attributes published in less-than-ideal ways.

Given the extensive variability in attribute names and their usage, ensuring comprehensive extraction of necessary information while avoiding erroneous interpretation from irrelevant data is challenging for rule-based methods. This underscores the value of using LLMs for metadata organization, as they are able to flexibly interpret texts based on contexts even with such complex and inconsistent descriptions.

### Creation of the Gold Standard Dataset for Cell Line Extraction

To evaluate the performance of the cell line extraction task, we created a gold standard dataset (https://doi.org/10.5281/zenodo.14643285). This dataset includes 300 samples derived from ChIP-seq experiments and 300 samples derived from ATAC-seq experiments. To ensure fairness in sample selection, only samples originating from different projects and assigned to distinct classifications in ChIP-Atlas were included in each set.

For samples considered to represent cell lines, the corresponding Cellosaurus terms were assigned and defined as the correct mappings. The final gold standard dataset was validated through manual inspection by two developmental biology experts and two bioinformatics experts, ensuring a high level of reliability.

Out of the selected samples, the number identified as representing cell lines was 183 for the ChIP-seq set and 139 for the ATAC-seq set. Among these, 17 and 12 samples, respectively, were confirmed to be cell lines but lacked corresponding terms in Cellosaurus. We recognize that such samples may include instances where submitters have assigned unique names to cell lines.

### Comparison of Existing Methods and LLM-Assisted Approaches Using the Gold Standard Dataset

Using the constructed gold standard dataset, we evaluated whether concept extraction using an LLM improves upon existing methods. Cell line names were extracted using the Llama 3.1 70B model, following the workflow illustrated in Fig. 1.

We compared the LLM-assisted pipeline with the MetaSRA pipeline using the metrics described in the Method section (Table 4). The gold standard dataset includes samples from both ChIP-seq and ATAC-seq experiments to evaluate the effect of gene names on the task, but the LLM did not mistakenly extract gene names as cell line names in any case, and no significant differences were observed between the two types of samples (Supplementary Table 1). Conventional methods achieved high accuracy in mapping cell line samples to ontology terms by restricting the attribute names used. This conservative strategy also enabled a high probability of correctly identifying non-cell line samples. However, this approach came at the cost of cell line coverage, resulting in many actual cell line samples being left unmapped. In contrast, the LLM-assisted method achieved high coverage in ontology mapping for cell line samples without compromising accuracy by selecting the most appropriate strings from all available attributes. At the same time, the samples that remained unmapped to cell line terms were more likely to be genuinely non-cell line samples. This underscores the efficacy of LLM-based methods in enhancing the quality of automatic metadata curation.

**Table 4.** The evaluation of the cell line extraction task by conventional and suggesting methods.

| Pipeline | Cell line accuracy | Cell line coverage | Non-cell line precision | Non-cell line recall |
| --- | --- | --- | --- | --- |
| MetaSRA | 0.903 | 0.721 | 0.782 | 0.937 |
| LLM-assisted | 0.923 | 0.930 | 0.940 | 0.934 |

Table 5 categorizes and quantifies the errors made by the LLM for samples where it failed to produce the correct output. Note that this categorization does not necessarily cover all possible errors that may occur in future executions. The most common error involved cases where the input mentioned a cell line, but the sample represented a derivative of that cell line rather than the cell line itself. The LLM were likely to incorrectly identify these as cell lines. In other cases, the LLM overlooked cell line names in input metadata. Among the eight samples where this failure occurred, six did not have attributes including either “cell line” or “cell type” in their keys. While the remaining two samples had a “cell type” attribute, the strings that should have been extracted were relatively short (”H1” and “JK1”).

**Table 5.** The categorization of the errors made by the LLM.

|  |  |
| --- | --- |
| Derivation: The sample is not the cell line itself, but is derived from the cell line | 12 |
| Overlook: The cell line name is overlooked by LLM | 8 |
| Non-canonical name: The cell line name is not canonical and not correspond to any Cellosaurus terms | 8 |
| Selection failure: LLM failed to select the correct mapping from multiple candidates | 6 |
| Wrong extraction: extracted string does not represent the cell line name | 5 |
| Ontology insufficiency: The term in Cellosaurus matching the string does not actually represent the cell line | 2 |
| **Total** | **41** |

Ontology mapping of the extracted strings resulted in multiple candidate Cellosaurus terms for 26 samples. The LLM was tasked with selecting the most likely cell line from the candidates (Prompt 2).

The prompt instructed the LLM to withhold judgment if the information provided in the BioSample metadata was insufficient to narrow down the candidates.

Of the 26 samples, 8 were judged as incorrect for reasons other than “Selection failure” in Table 5. Among the remaining 18 samples, the LLM correctly selected the appropriate cell line for 11 samples and appropriately withheld judgment for 1 sample. In 4 cases, it incorrectly selected a single cell line when it should have withheld judgment due to insufficient information. In 2 cases, it incorrectly withheld judgment when it was expected to identify the appropriate cell line based on the BioSample descriptions. Taken together, for samples where a decision was feasible, the LLM achieved 11 correct answers out of 13. These results suggest that the LLM can be effectively employed for tasks requiring the differentiation of identically named cell lines.

These findings suggest that while some challenges remain in disambiguating sample context and ensuring comprehensive extraction, the LLM-assisted approach can substantially improve performance.

### Evaluation of A Potential Application Extraction of Experimentally Altered Gene Names and Techniques

Based on the evaluation results of the cell line extraction task, we concluded that concept extraction using LLMs can be performed at a practical level for biological experimental factors. With this in mind, we aimed to enhance the utility of existing applications by applying similar methods to concepts not yet covered by manual curation.

We attempted to extract information about genes whose expression was experimentally modulated from the metadata of experimental samples collected by ChIP-Atlas. As described in the Methods section, we used a total of 3,723 samples, consisting of 1,752 ATAC-seq samples and 1,971 ChIP-seq samples. Using Prompt 3 from the Methods section for extraction, at least one gene was identified in 600 out of the 3,723 samples. These results were manually evaluated for correctness, assessing the accuracy of gene names and method names separately. We excluded samples for which a single correct answer is not clearly defined in the prompt. Examples of such cases include samples mentioning fusion genes, where it was unclear whether individual gene names within a fusion should be extracted separately or combined using notation like hyphens or double-colon (::). Other examples are samples with a mutated gene. In some cases the LLM output “mutation” as the method name, even if the input lacked this word, while in other cases it extracted terms like “K36M” exactly as described. Both were deemed reasonable in practice but we excluded them from evaluation because the prompt does not define which is correct.

Out of the 600 extractions, 579 cases were evaluable for both gene names and method names and the accuracy rate was 0.803. When evaluated separately, the accuracy for gene names was 0.916, and the accuracy for method names was 0.847.

The extraction results included 459 unique gene names. Using the HGNC Multi-symbol checker, 396 of these were mapped to one or more HGNC IDs. Among these, 32 were assigned multiple IDs and could not be uniquely resolved. Although gene symbols defined by HGNC are unique across all human genes, they are not always unique when synonyms are included. The information described in BioSample alone is typically insufficient to distinguish between them, representing a challenge for future work. Meanwhile, 63 gene names could not be mapped to any corresponding ID. These cases included scenarios where genes from non-human organisms, such as GFP, had been introduced, as well as cases where common names employing Greek letters not recognized by HGNC nomenclature were used.

Coverage was not evaluated in this study due to the absence of pre-existing manually curated results. However, the results of accuracy evaluation demonstrate the potential for LLM-assisted extraction to significantly reduce the effort required by database users during sample searches.

Extraction results judged as incorrect often involved complex descriptions. Table 6 provides examples of such cases. These include, for example, a sample where only the name of an inhibitor was mentioned and additional information was required to determine the affected gene. Another example is a sample where only the transduced gene carries an amino acid substitution mutation. This presented challenges, as describing them comprehensively requires defining a complex schema.

**Table 6.**
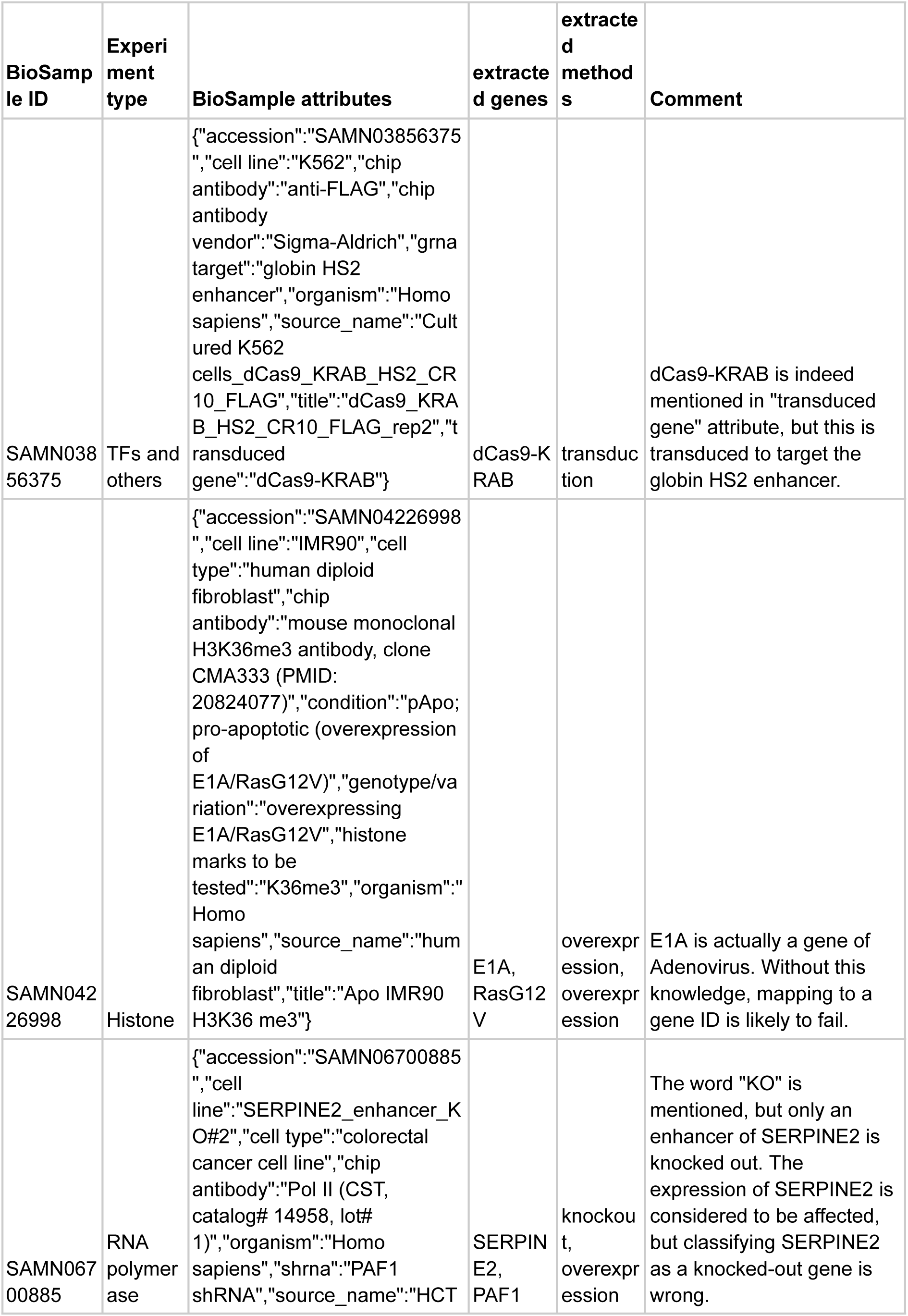
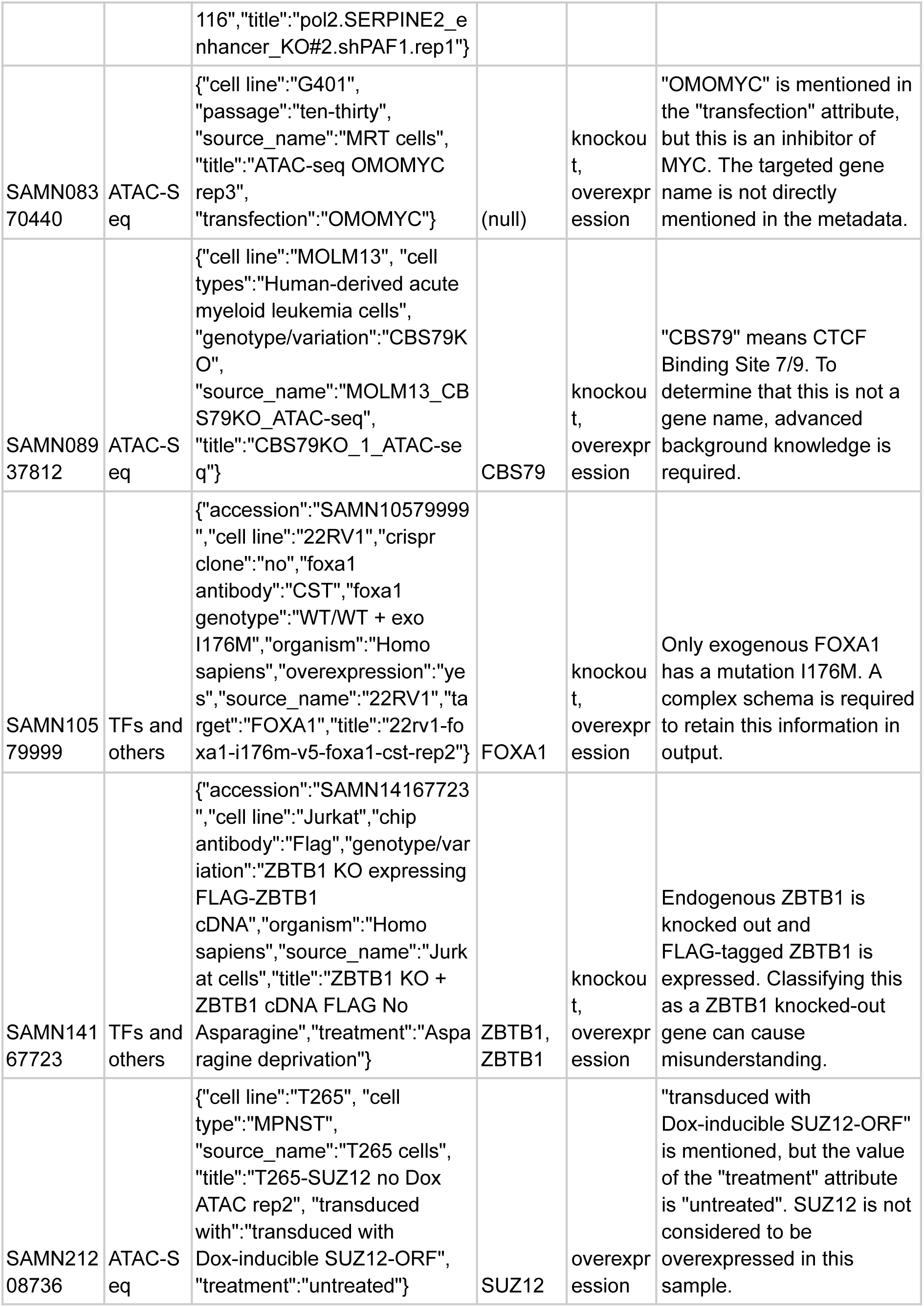
Examples of BioSample records with attributes that are difficult to describe in a simple schema.

Designing prompts to account for every possible case is impractical. Instead, each application must find an appropriate balance of accuracy and coverage, based on their specific requirements.

## Discussion

### Outcomes

In this study, we quantitatively confirmed the effectiveness of LLMs for extracting cell line names—a concept that has already been subject to manual curation—using samples covered by ChIP-Atlas. We further applied the same approach to the extraction of gene names that have been experimentally modulated. When searching ChIP-seq experimental data using simple string matching, the gene names retrieved often represent a mix of targets used in ChIP experiments and targets subjected to manipulations such as knockouts. The current version of ChIP-Atlas has allowed filtering of experiments only by the type of cells or tissues used. Within the same classification, however, there can be samples where expressions of some genes have been experimentally modulated. By curating such information with LLMs, users could exclude samples involving KO or KD to reduce noise in their analysis and focus on more relevant results.

Although LLMs are expected to assist in correcting low-quality metadata generated by humans, efforts to prevent the creation of such low-quality metadata in the first place remain essential. For example, NCBI has been working to improve the quality of submitted metadata by introducing additional constraints that metadata must meet upon submission to BioSample and by enhancing documentation [20]. Tools such as CEDAR [21] are also available to assist in metadata creation. Data submitters should take advantage of these support systems and recognize that submitting data to public repositories is intended to enable data reuse. While the data submitters should ensure that their metadata is properly described, we also understand the errors and mistakes can be published unintentionally. The outcomes of this study can handle those errors to rescue those published data for further use.

Under the evaluation environment used in this study, the LLM could process approximately 400 samples per hour. The total number of epigenomics experiments included in ChIP-Atlas is approximately 430,000, which can be processed within a practical timeframe, enabling the benefits of LLM-based curation to be directly translated into user utility. Still, it should be noted that the total number of records in BioSample exceeds 40 million, and addressing this larger scale would require additional pre-processing or advancements in model performance.

Another key contribution of this research is demonstrating the utility of Llama 3, a locally deployable model. While many studies rely on commercial models such as GPT by OpenAI [22], our adoption of a local model ensures transparency and sustainability, avoiding dependence on specific vendors. Additionally, for large-scale and continuous data processing, relying on paid services could raise sustainability concerns. Moreover, using a local model like Llama 3 makes it feasible to apply similar methods to sensitive data, such as electronic health records, where privacy is paramount. Also, our approach is more flexible than methods that depend on fixed schemas like those employed by the CEDAR group [16], allowing term extraction from arbitrary text rather than requiring adherence to predefined structures.

### Limitations

Despite these advances, several challenges remain unresolved:

1. **Complex metadata descriptions**: Experimental sample metadata can be intricate, making it difficult to represent some cases with the simple schema used in this study. For instance:

○ **Differentiated cell types**: When describing samples of cells differentiated from a specific cell line, ideally, both the original cell line and the differentiated cell type should be recorded.
○ **Fusion proteins**: For gene name extraction, mapping to NCBI Gene IDs is complicated because NCBI Gene lacks entries for fusion genes. This necessitates using individual gene IDs and designing schemas that convey the information about the fusion gene as a whole, not just its components.
2. While schema design and prompt engineering can partially address these issues, complete automation remains challenging.
3. **Limits of prompt engineering**: While improvements in model performance may yield better results for the same prompts, predicting the extent of these improvements is difficult.
4. **Computational constraints**: Processing the entirety of BioSample would require extensive computational resources, time, and energy. These constraints necessitate careful consideration of the practical scope and application of LLM-based approaches for each specific task.

In light of these limitations, achieving fully comprehensive results with current LLMs may not be feasible for all tasks. Instead, it is essential to define appropriate use cases and balance expectations based on available resources and application needs.

### Future Directions

The rapid advancement of LLMs holds significant promise for tasks like experimental metadata curation. As more powerful models become available, we anticipate further improvements in performance. As shown by the usefulness of ChIP-Atlas’s manual curation results in this study, human curation remains valuable for providing near-complete curation and for evaluating the effectiveness of automated methods. Still, LLMs are poised to significantly reduce the workload of human curators.

This study represents an initial step in this direction, laying the groundwork for future applications and refinements. With continued development, LLM-based methods are expected to play a critical role in bridging the gap between large-scale metadata and efficient, accurate curation processes.

## Notes

### Competing Interest Statement

The authors have declared no competing interest.

## References

1. Katz K, Shutov O, Lapoint R, Kimelman M, Brister JR, O’Sullivan C. The Sequence Read Archive: a decade more of explosive growth. Nucleic Acids Res. 2022 Jan 7;50(D1):D387–90.

2. Clough E, Barrett T. The Gene Expression Omnibus database. Methods Mol Biol Clifton NJ. 2016;1418:93–110.

3. Ziemann M, Kaspi A, El-Osta A. Digital expression explorer 2: a repository of uniformly processed RNA sequencing data. GigaScience. 2019 Apr 3;8(4):giz022.

4. Mahi NA, Najafabadi MF, Pilarczyk M, Kouril M, Medvedovic M. GREIN: An Interactive Web Platform for Re-analyzing GEO RNA-seq Data. Sci Rep. 2019 May 20;9(1):7580.

5. Zou Z, Ohta T, Oki S. ChIP-Atlas 3.0: a data-mining suite to explore chromosome architecture together with large-scale regulome data. Nucleic Acids Res. 2024 Jul 5;52(W1):W45–53.

6. Barrett T, Clark K, Gevorgyan R, Gorelenkov V, Gribov E, Karsch-Mizrachi I, et al. BioProject and BioSample databases at NCBI: facilitating capture and organization of metadata. Nucleic Acids Res. 2012 Jan 1;40(D1):D57–63.

7. Gonçalves RS, Musen MA. The variable quality of metadata about biological samples used in biomedical experiments. Sci Data. 2019 Feb 19;6(1):190021.

8. Mungall CJ, Torniai C, Gkoutos GV, Lewis SE, Haendel MA. Uberon, an integrative multi-species anatomy ontology. Genome Biol. 2012 Jan 31;13(1):R5.

9. Bard J, Rhee SY, Ashburner M. An ontology for cell types. Genome Biol. 2005;6(2):R21.

10. Schriml LM, Munro JB, Schor M, Olley D, McCracken C, Felix V, et al. The Human Disease Ontology 2022 update. Nucleic Acids Res. 2021 Nov 10;50(D1):D1255–61.

11. Bernstein MN, Doan A, Dewey CN. MetaSRA: normalized human sample-specific metadata for the Sequence Read Archive. Bioinformatics. 2017 Sep 15;33(18):2914–23.

12. Bairoch A. The Cellosaurus, a Cell-Line Knowledge Resource. J Biomol Tech JBT. 2018 Jul;29(2):25–38.

13. Malone J, Holloway E, Adamusiak T, Kapushesky M, Zheng J, Kolesnikov N, et al. Modeling sample variables with an Experimental Factor Ontology. Bioinformatics. 2010 Apr 15;26(8):1112–8.

14. Klie A, Tsui BY, Mollah S, Skola D, Dow M, Hsu CN, et al. Increasing metadata coverage of SRA BioSample entries using deep learning–based named entity recognition. Database. 2021 Sep 29;2021:baab021.

15. Dagdelen J, Dunn A, Lee S, Walker N, Rosen AS, Ceder G, et al. Structured information extraction from scientific text with large language models. Nat Commun. 2024 Feb 15;15(1):1418.

16. Sundaram SS, Solomon B, Khatri A, Laumas A, Khatri P, Musen MA. Use of a Structured Knowledge Base Enhances Metadata Curation by Large Language Models [Internet]. arXiv; 2024 [cited 2024 Jun 19]. Available from: http://arxiv.org/abs/2404.05893

17. Ollama [Internet]. [cited 2024 Dec 17]. Available from: https://ollama.com

18. Introducing Llama 3.1: Our most capable models to date [Internet]. Meta AI. [cited 2024 Dec 17]. Available from: https://ai.meta.com/blog/meta-llama-3-1/

19. Kojima T, Gu SS, Reid M, Matsuo Y, Iwasawa Y. Large Language Models are Zero-Shot Reasoners [Internet]. arXiv; 2023 [cited 2024 Dec 2]. Available from: http://arxiv.org/abs/2205.11916

20. Staff N. Upcoming Changes to NCBI’s BioSample Database [Internet]. NCBI Insights. 2024 [cited 2025 Jan 23]. Available from: https://ncbiinsights.ncbi.nlm.nih.gov/2024/10/23/changes-ncbis-biosample-database/

21. Gonçalves RS, O’Connor MJ, Martínez-Romero M, Egyedi AL, Willrett D, Graybeal J, et al. The CEDAR Workbench: An Ontology-Assisted Environment for Authoring Metadata that Describe Scientific Experiments. Semantic Web--ISWC Int Semantic Web Conf Proc Int Semantic Web Conf. 2017 Oct;10588:103–10.

22. ChatGPT [Internet]. [cited 2023 Dec 27]. Available from: https://chat.openai.com

