## Supplementary for "Extraction of biological terms using large language models enhances the usability of metadata in the BioSample database"

### DNA-seq of Human: CRC patient HO110-2011D

Related information
BioProject
SRA
Taxonomy

Recent activity

| | |
|---|---|
| Identifiers | BioSample: SAMN18236550; Sample name: HO110-2011D; SRA: SRS8421728 |
| Organism | Homo sapiens (human)<br>cellular organisms; Eukaryota; Opisthokonta; Metazoa; Eumetazoa; Bilateria; Deuterostomia; Chordata; Craniata; Vertebrata; Gnathostomata; Teleostomi; Euteleostomi; Sarcopterygii; Dipnotetrapodomorpha; Tetrapoda; Amniota; Mammalia; Theria; Eutheria; Boreoeutheria; Euarchontoglires; Primates; Haplorrhini; Simiiformes; Catarrhini; Hominoidea; Hominidae; Homininae; Homo |
| Package | Human; version 1.0 |

Attributes

**isolate**
**age**
**biomaterial provider**
**sex**
**tissue**
**sample type**
**Treatment date**
**Pathology No:**
**Histological type:1.adeno ca 2.mucinous ca(mucin>50%) 3.signet call ca 4.squanous ca 5.adenosquamous ca 6.small cell c**
**grade of differentiation:1.well 2.moderately 3.poorly 4.undifferentiated 5 no data**
**other pathological parameters:1.Blood Vascular Invasion 2.Lymphatic Vessel Invasion 3.perineural invasion 4.isolated can**
**Mucinous component %**
**No of metastatic nodes/No of total sampling nodes: /**
**apical nod or root nodes metastasis 0: No 1: yes**
**T(is-4)**
**N(1-3)**
**M(0-9)**
**Shape of advanced tumor Borrmann type 1: type I 2:type II 3:type III 4:type IV**
**Size of tumor(according to the pathology report) x x cm**
**Date of recurrence found**

### Microbe sample from Oenococcus oeni

BioProject
SRA
Taxonomy

Recent activity

| | |
|---|---|
| Identifiers | BioSample: SAMN06840936; Sample name: 160837B; SRA: SRS2150895 |
| Organism | Oenococcus oeni<br>cellular organisms; Bacteria; Bacillati; Bacillota; Bacilli; Lactobacillales; Lactobacillaceae; Oenococcus |
| Package | Microbe; version 1.0 |

Attributes

| | |
|---|---|
| **host** | not applicable |
| **isolation source** | wine |
| **collection date** | 2000-06-01 |
| **geographic location** | China:Shandong |
| **latitude and longitude** | 37.32 N 121.24 E |
| **strain** | SD-2a |
| **isolate** | not applicable |
| **sample type** | cell culture |
| **altitude** | missing |
| **biomaterial provider** | missing |
| **collected by** | missing |
| **culture collection** | missing |
| **depth** | missing |
| **broad-scale environmental context** | missing |
| **genotype** | missing |
| **3** | pH3.0_1h_1 |
| **5** | pH3.0_1h_2 |
| **14** | pH3.0_1h_3 |
| **9** | pH4.8_0h_1 |
| **8** | pH4.8_0h_2 |
| **15** | pH4.8_0h_3 |
| **13** | pH4.8_1h_1 |
| **12** | pH4.8_1h_2 |
| **10** | pH4.8_1h_3 |
| **160837B-3** | pH3.0_1h_1 |
| **160837B-5** | pH3.0_1h_2 |
| **160837B-14** | pH3.0_1h_3 |
| **160837B-9** | pH4.8_0h_1 |

Supplementary figure 1. Screenshots of NCBI BioSample webpages for records with attributes described in less-than-ideal ways. (A) SAMN18236550. This has attributes whose name defines option numbers and their corresponding values, resulting in extreme length. Since the web interface of

NCBI BioSample does not assume such long attribute names, it cannot display full-length attributes and their values. (B) SAMN06840936. This has uninterpretable attributes like “3” or “160837B-3”. Inferring from their values, they may mean pH and time for certain treatment, but we cannot say which condition was used for this specific sample.

Supplementary Table 1. The evaluation of the cell line extraction task per each experiment type by conventional and suggesting methods.

| **Experiment type** | **Pipeline** | **Cell line accuracy** | **Cell line coverage** | **Non-cell line precision** | **Non-cell line recall** |
|---|---|---|---|---|---|
| ATAC | MetaSRA | 0.910714 | 0.856383 | 0.784615 | 0.947059 |
| ChIP | MetaSRA | 0.896825 | 0.701149 | 0.672619 | 0.924242 |
| ATAC | LLM-assisted | 0.902985 | 0.957831 | 0.930769 | 0.935294 |
| ChIP | LLM-assisted | 0.939759 | 0.91791 | 0.928571 | 0.931818 |
